# Long-read PacBio HiFi sequencing resolves the genomic integration site and structure of the Nestin-CreERT2^KEisc/J^ transgene

**DOI:** 10.64898/2026.10.01.755969

**Authors:** Alycia M Charest, Alain Charest

## Abstract

Random integration transgenesis has been foundational to modern mouse genetics, generating thousands of lines that continue to serve as important tools in biomedical research. Yet for many of these models, the genomic integration site and associated genomic alterations remain unknown, complicating zygosity determination and potentially obscuring disruption of endogenous loci. Mice expressing inducible CreERT2 under rat *Nes* regulatory elements are widely used for *in vivo* lineage tracing of neural stem and progenitor cells, yet the genomic integration site of the Tg(Nescre/ERT2)KEisc/J transgene has remained undefined. Here, we used PacBio HiFi whole-genome sequencing to resolve its genomic locus and molecular architecture. We reconstructed the complete 13.7-kb transgene and mapped its integration to chromosome 6 within intron 1 of *Kcnd2*, where insertion was accompanied by deletion of 17.4 kb of endogenous genomic DNA. Identification of the integration junctions further enabled development of an allele-specific PCR assay for direct zygosity determination. Beyond resolving the molecular architecture of a widely used CreERT2 mouse line, this study demonstrates the utility of long-read HiFi sequencing for revisiting legacy transgenic models, uncovering otherwise hidden genomic alterations while enabling precise molecular genotyping.

**Statement of Significance:** identifying genomic integration sites of foreign DNA in transgenic animals can be a laborious endeavor. The advent of long-read genomic sequencing (PacBio HiFi) facilitates such identification and allows for development of accurate genotyping of the transgene DNA. The pipeline described in this manuscript can be readily adapted to any transgene integration search.

## Introduction

The development of transgenic mice through pronuclear injection of exogenous DNA into fertilized eggs fundamentally transformed mammalian genetics by enabling the study of gene function in the context of the intact animal. In conventional pronuclear transgenesis, injected DNA integrates into the genome at an essentially random location, allowing regulatory sequences to be used to direct transgene expression to defined tissues and cell populations (1,2). The subsequent development of the Cre/loxP recombination system greatly expanded the utility of mouse genetics by permitting conditional modification of endogenous genes. When Cre driver strains are combined with alleles containing strategically positioned loxP sites, genes can be deleted, activated, or otherwise modified selectively within Cre-expressing cells, circumventing developmental lethality and other limitations associated with conventional germline mutations (3).

An important further advance was the development of ligand-inducible Cre recombinases, particularly CreER and CreERT2. Fusion of Cre recombinase to a modified estrogen receptor ligand-binding domain renders Cre activity dependent upon administration of tamoxifen or its active metabolites. In the absence of ligand, CreER/CreERT2 is largely excluded from the nucleus, whereas ligand administration promotes nuclear translocation and subsequent recombination of loxP-flanked sequences (4,5). The combination of tissue-specific regulatory elements with CreERT2 therefore provides both spatial and temporal control of genetic modification and has become a powerful approach for lineage tracing, conditional gene deletion, gene activation, and functional studies during embryonic development and in adult tissues.

The recognition that neural stem and progenitor cells persist in the adult mammalian brain fundamentally changed concepts of adult brain plasticity. In rodents, adult neurogenesis occurs prominently in the ventricular-subventricular zone (V-SVZ), whose progenitors generate neurons that migrate toward the olfactory bulb through the rostral migratory stream, and in the subgranular zone of the hippocampal dentate gyrus, where newly generated granule neurons integrate into existing neuronal circuits. These discoveries created a need for genetic systems capable of identifying through lineage tracing neural stem and progenitor cells and manipulating them in a temporally controlled manner. Nestin, an intermediate filament protein expressed prominently in neural stem and progenitor populations, became one of the most widely used regulatory loci for this purpose.

In fact, up to 9 independently generated Nestin-CreERT2 transgenic mouse strains have been developed using regulatory elements derived from the rat *Nestin* (*Nes*) locus (6). Importantly, these strains were generated using different *Nes* regulatory constructs and independent genomic integration events and therefore represent genetically distinct transgenic lines rather than interchangeable versions of a single model. One widely used strain, Tg(Nes-cre/ERT2)KEisc (JAX stock #016261), was generated by Lagace and colleagues to inducibly label and genetically manipulate nestin-expressing neural stem cells and their progeny in the adult brain (7). In this strain, CreERT2 was cloned into a construct that contains approximately 5.8 kb of the rat *Nes* promoter together with downstream *Nes* genomic sequences encompassing portions of exons 1 and 4, complete exons 2 and 3, and the intervening introns (7). Tamoxifen administration induces recombination in neural stem and progenitor populations within the adult V-SVZ and subgranular zone (6,7).

Additional Nestin-CreERT2 lines have been independently generated using different configurations of *Nes* regulatory sequences and a direct head-to-head comparison of three of these strains has demonstrated substantial differences in the distribution, specificity, and efficiency of Cre-mediated recombination (6). This lack of consistency may result from differences in the regulatory constructs themselves as well as from the distinct genomic environments into which the transgenes integrated. Consequently, findings obtained using one Nestin-CreERT2 strain cannot necessarily be assumed to apply to another, emphasizing the importance of accurately defining the molecular identity of each individual transgenic line.

A fundamental feature of classical pronuclear transgenesis is that the genomic location of transgene integration is not predetermined. The genomic environment surrounding an integration site can influence transgene expression, while insertion itself can potentially disrupt endogenous genes or regulatory sequences. Knowledge of the precise integration site therefore provides an important component of the molecular characterization of a transgenic strain. In addition to identifying potential effects on the endogenous genome, determination of the integration locus can help distinguish independently generated strains, provide insight into differences in transgene expression among lines, and enable the development of integration-junction-specific genotyping assays.

Determining transgene integration sites using conventional short-read whole-genome sequencing approaches can be challenging because of difficulties resolving insertion boundaries within repetitive or structurally complex regions. Long-read sequencing provides a powerful alternative since individual reads can extend from transgene-derived sequences directly into the adjacent endogenous genome. PacBio circular consensus sequencing (CCS), or HiFi sequencing, combines long read lengths with high per-base accuracy and is therefore particularly well suited for identifying and resolving such transgene-genome boundaries (8).

Here, we used PacBio HiFi whole-genome sequencing to determine the genomic integration site of the widely used Tg(Nes-cre/ERT2)KEisc Nestin-CreERT2 mouse strain. By identifying long reads containing Nestin-CreERT2 sequences and mapping sequences associated with the transgene to the mouse reference genome, we identified the chromosomal integration locus and defined its genomic boundary, providing a molecular characterization of a Nestin-CreERT2 strain that has been widely used for inducible genetic manipulation of neural stem and progenitor cells.

## Materials and Methods

### Mouse maintenance

All mouse procedures were carried out in accordance with Beth Israel Deaconess Medical Center recommendations for care and use of animals and were maintained and handled under protocols approved by Institutional Animal Care and Use Committee (IACUC).

### High-molecular-weight DNA extraction and PacBio HiFi whole-genome sequencing

Spleens were harvested from mice heterozygous for the germline Tg(Nes-cre/ERT2)KEisc transgene [C57BL/6-Tg(Nes-cre/ERT2)KEisc/J; The Jackson Laboratory, stock no. 016261] and mechanically dissociated to obtain approximately 1 × 10^7 cells for high-molecular-weight (HMW) genomic DNA extraction. HMW genomic DNA was isolated using PacBio Nanobind PanDNA kit [Pacific Biosciences, #103-260-300] according to the manufacturer’s instructions. Purified HMW genomic DNA was submitted to Azenta Life Sciences for PacBio HiFi whole-genome sequencing. Genomic DNA was sheared using a Megaruptor 3 system, and PacBio SMRTbell libraries were prepared with bead-based size selection according to the manufacturer’s protocol. Libraries were sequenced on a PacBio Revio system (Pacific Biosciences), and HiFi reads were generated using SMRT Link v25.1 and provided in BAM format. Sequencing generated 6,961,154 HiFi reads comprising 102.03 Gb of sequence, with a mean HiFi read length of 14.66 kb. Based on an approximately 2.7-Gb haploid mouse genome, the total sequence yield corresponded to approximately 38× theoretical whole-genome coverage.

### Data processing and alignment

PacBio HiFi reads were analyzed using command-line sequence analysis tools on a local computing workstation. Sequence manipulation and read extraction were performed using SeqKit v2.13.0 and SAMtools v1.24, and sequence alignments were performed using minimap2 v2.31-r1302. PacBio HiFi reads were provided in BAM format and converted to FASTQ format for downstream sequence analysis. To identify HiFi reads containing sequences derived from the Nestin-CreERT2 transgene, the complete HiFi read dataset was screened against the known CreERT2 coding sequence using minimap2 v2.31-r1302. CreERT2-containing reads were extracted using SeqKit v2.13.0 and SAMtools v1.24, yielding 1,091 unique full-length HiFi reads. These reads were subsequently aligned and manually assembled based on overlapping sequence to reconstruct transgene-containing sequence contigs. Resulting contigs were examined for CreERT2 and adjacent transgene sequences, and sequences extending beyond the transgene-containing regions were aligned to the GRCm39 mouse reference genome to identify endogenous genomic sequences flanking the transgene insertion. Mapping of these flanking sequences localized the Nestin-CreERT2 integration site to chromosome 6 (NC_000072.7) and identified genomic sequences flanking both sides of the insertion.

Comparison of the assembled transgene-containing sequence with the corresponding chromosome 6 reference sequence was then used to define the approximate transgene-genome junctions and reconstruct the integrated Nestin-CreERT2 sequence. This analysis generated a 13,714-bp reconstructed Nestin-CreERT2 transgene sequence, which was subsequently used as a reference for read-level validation of the transgene structure and integration site. To independently evaluate read support for the identified locus, a 100-kb GRCm39 reference interval spanning chr6:21,200,000– 21,300,000 was extracted and used as a focused genomic reference. The complete PacBio HiFi dataset was aligned independently against this chromosome 6 reference sequence and the reconstructed 13,714-bp Nestin-CreERT2 sequence using minimap2 with the HiFi read preset (-x map-hifi). For recruitment of reads mapping to the chromosome 6 interval, alignments of ≥2,000 bp with ≥95% sequence identity were retained. A total of 265 HiFi reads satisfied these criteria. Read identifiers from the chromosome 6 and transgene alignments were then compared to identify reads containing sequence support for both the endogenous chromosome 6 locus and the Nestin-CreERT2 transgene. This analysis identified 25 PacBio HiFi reads supporting the transgene integration locus. Individual reads were examined with respect to their orientation and alignment to the endogenous chromosome 6 sequence and the reconstructed Nestin-CreERT2 sequence. Fifteen reads supported the left transgene-genome junction and converged near GRCm39 chromosome 6 position 21,251,189, whereas ten reads supporting the right junction converged near position 21,268,582, with one read differing by approximately 10 bp. Junction sequences were subsequently inspected and manually refined, defining the integration boundaries at approximately chr6:21,251,200 and chr6:21,268,569 (GRCm39). These coordinates indicate replacement of approximately 17.4 kb of endogenous chromosome 6 sequence at the transgene integration site.

### PCR genotyping

Ear-punch tissue samples were processed for direct PCR DNA isolation using a commercially available Mouse Direct PCR kit (Selleckchem) according to the manufacturer’s instructions. Briefly, ear-punch tissues were submerged in 100 µL of lysis buffer containing 2 μL of Protease Plus and incubated at 55°C for 1 h, followed by incubation at 95°C for 10 min. The resulting DNA lysates were diluted fivefold in ddH_2_O, and 5 µL of diluted DNA was used as template in a 20-µL PCR reaction containing 0.25 µM of each primer along with 2x PCR mix containing the polymerase. PCR amplification was performed using the following cycling conditions: initial denaturation at 94°C for 5 min; 37 cycles of denaturation at 94°C for 30 s, annealing at 55°C for 30 s, and extension at 72°C for 30 s; followed by a final extension at 72°C for 10 min. Reactions were subsequently held at 10°C. PCR products were resolved by electrophoresis on 2.5% agarose gels in TAE buffer, stained with ethidium bromide, and visualized and photographed under UV illumination. Primer sequences and expected PCR products are shown in Fig. 2.

### Data availability

The PacBio HiFi whole-genome sequencing data generated in this study have been deposited in the NCBI Sequence Read Archive (SRA) under BioProject accession PRJNA1533531 and SRA accession SRR40804357. The reconstructed 13,714-bp Tg(Nes-cre/ERT2)KEisc transgene sequence has been deposited in GenBank under accession QB046940. Additional data supporting the findings of this study are available from the corresponding author upon request.

## Results

### Mapping of the Nestin-CreERT2 transgene integration site and reconstruction of the transgene

To identify the genomic integration site of the Nestin-CreERT2 transgene, PacBio HiFi reads were initially screened using the known CreERT2 sequence, identifying 1,091 unique CreERT2-containing reads among the 6,961,154 CCS reads. These reads were used as an initial candidate pool to reconstruct the transgene and identify sequences extending beyond the transgene into flanking mouse genomic DNA. Alignment of these flanking sequences to the GRCm39 mouse reference genome localized the integration site to chromosome 6 cytogenetic band A2 (Fig. 1A), within the interval between approximately 21,240,000 and 21,280,000 bp (Fig. 1B). A total of 25 PacBio HiFi reads supported the reconstructed transgene-genome junctions, and six were tiled across the integration locus (Supplementary Fig. 1). Subsequent computational and manual sequence alignment refined the integration breakpoints to Chr6:21,251,200 and Chr6:21,268,569, defining a 17,369-bp deletion of the endogenous chromosome 6 sequence associated with insertion of the 13,714-bp Nestin-CreERT2 transgene.

**Figure 1.**
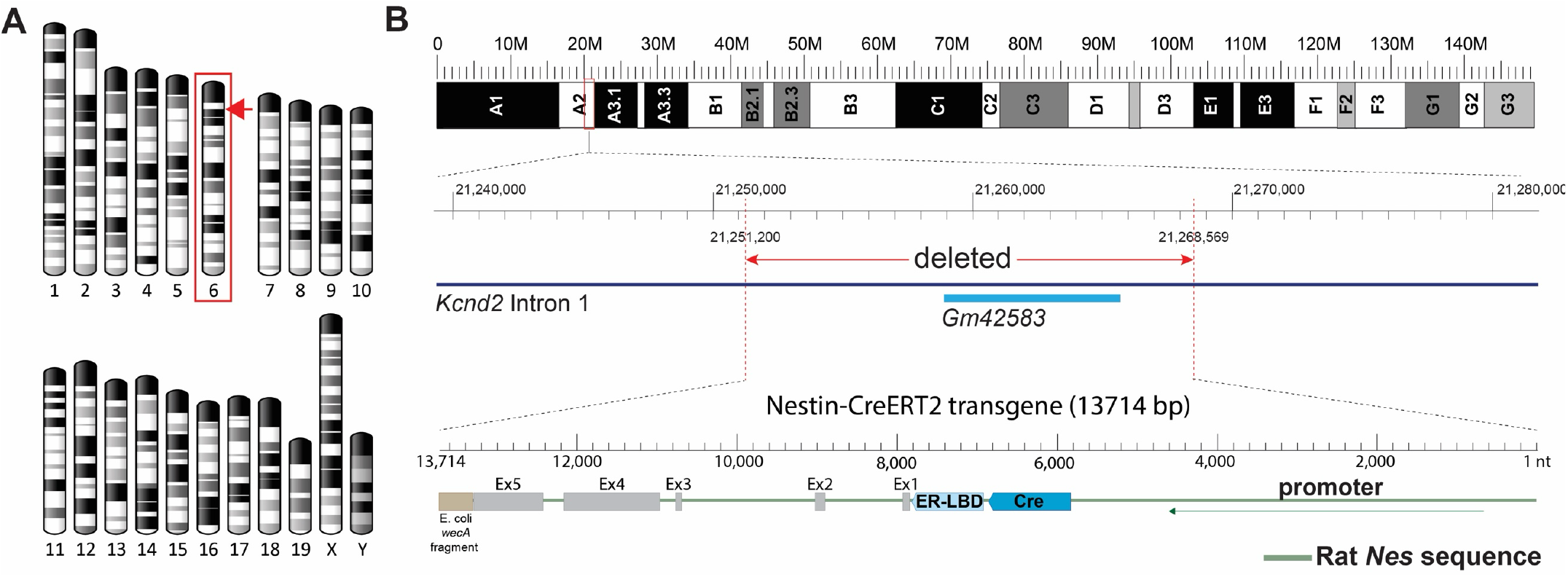
Genomic integration site and molecular structure of the Nestin-CreERT2 transgene. **(A)** Mouse chromosome idiograms showing the integration site within band A2 of chromosome 6 (arrow). **(B)** Genomic localization of the transgene integration site within intron 1 of *Kcnd2* on the GRCm39 mouse reference genome. Integration of the 13,714-bp Nestin-CreERT2 transgene is associated with deletion of 17,369 bp of endogenous genomic DNA between Chr6:21,251,200 and Chr6:21,268,569, encompassing the annotated *Gm42583* locus. The reconstructed transgene consists of rat *Nes* regulatory sequences driving CreERT2, followed by downstream rat *Nes* genomic sequences and a 409-bp fragment derived from the *E. coli wecA* locus. The transgene is integrated in the reverse transcriptional orientation relative to *Kcnd2*.

The complete 13,714-bp transgene sequence was reconstructed from the genomic sequencing data and deposited in GenBank (accession QB046940). The transgene contains 5,819 bp of the rat *Nes* 5’ region upstream of the first ATG translation initiation codon, which includes the experimentally defined transcription initiation site located 126 bp upstream of the ATG (9), and 123 bp of the native *Nes* 5′ untranslated region, thereby preserving essentially the entire native 5′ transcriptional context immediately preceding the translation initiation codon. This sequence is followed by the CreERT2 cDNA and rat *Nes* genomic sequences comprising exon 1, intron 1, exon 2, intron 2, exon 3, intron 3, exon 4, intron 4, and a portion of exon 5 (Fig. 1B). Immediately downstream of the rat *Nes* exon 5 sequence is a 409-bp fragment derived from the *Escherichia coli wecA* gene. The origin of this sequence within the transgene is unknown but may reflect incorporation of bacterial DNA during the original molecular cloning and construction of the transgene.

Integration of the transgene was accompanied by a 17.4-kb deletion within intron 1 of the *Kcnd2* gene (Fig. 1B). The transgene is oriented in the opposite transcriptional direction relative to *Kcnd2*. The deleted interval also encompasses Gm42583, an uncharacterized predicted transcript of unknown function (Ensembl biotype: TEC) (Fig. 1B). *Kcnd2* encodes Kv4.2, a voltage-gated potassium-channel subunit expressed in the central nervous system that contributes to the regulation of neuronal membrane excitability (10). Altered *Kcnd2*/Kv4.2 expression or function has been investigated in mouse models of several neurological and neuropsychiatric disorders, including epilepsy, autism spectrum disorders, and Alzheimer’s disease (11).

### Molecular validation of the integration site and development of allele-specific PCR genotyping

The candidate 5′ and 3′ transgene-genome junctions identified by computational analysis were further refined by manual local sequence alignment (Fig. 2A). Oligonucleotide primers were designed to flank each predicted transgene insertion breakpoint and the corresponding uninterrupted wild-type genomic sequence. Two three-primer PCR assays, independently targeting the 5′ and 3′ integration junctions, were developed to distinguish the transgenic allele from the endogenous wild-type allele (Fig. 2B).

**Figure 2.**
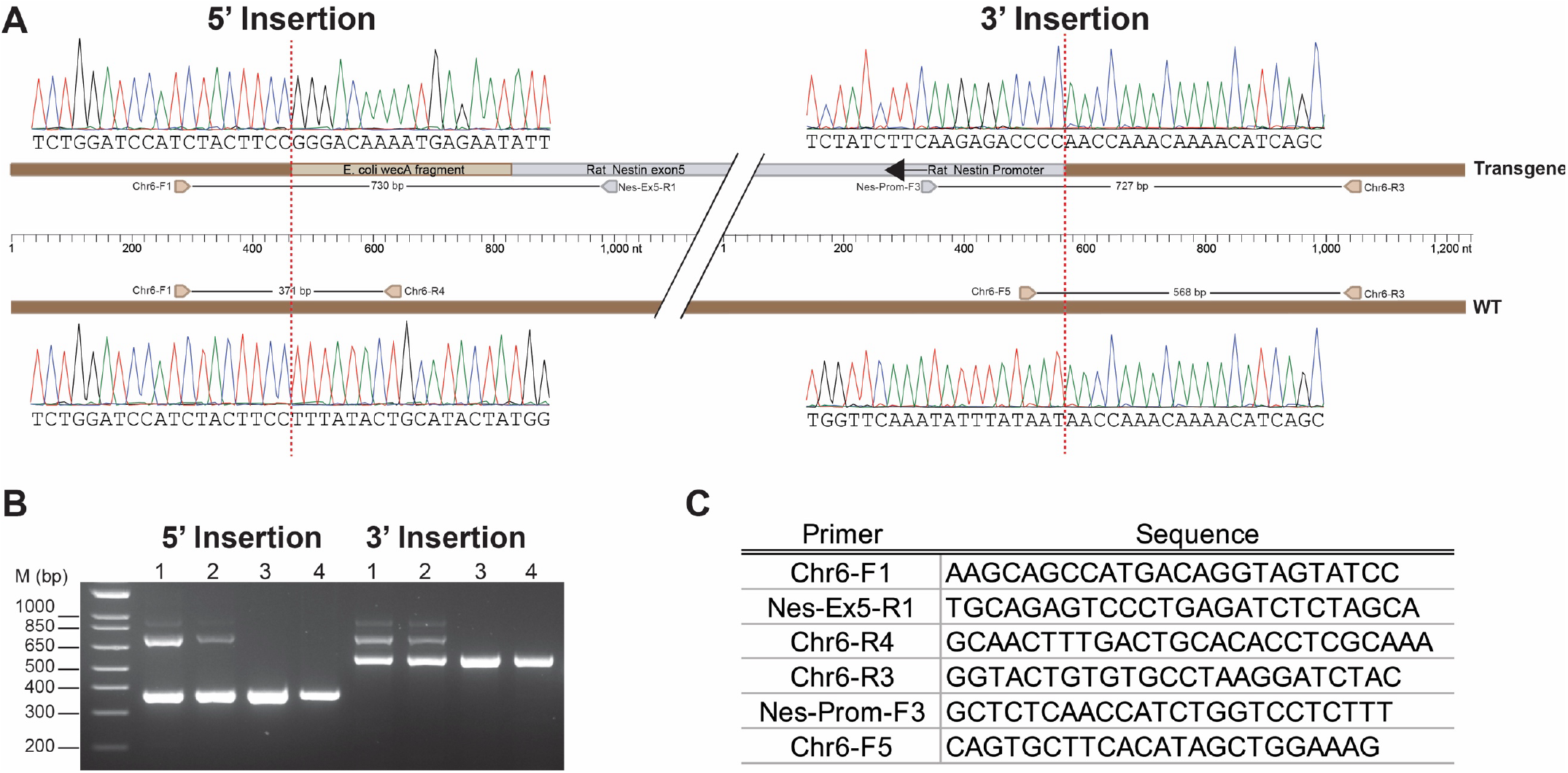
Nucleotide-level validation of the Nestin-CreERT2 transgene integration site and development of an allele-specific PCR genotyping assay. **(A)** Sanger sequencing chromatograms confirming the 5′ and 3′ transgene-genome junctions and the corresponding wild-type genomic sequences. Red dashed lines indicate the insertion breakpoints. Schematics show the location and orientation of primers used for allele-specific PCR genotyping and the expected PCR product sizes: 730 bp and 371 bp at the 5′ junction and 727 bp and 568 bp at the 3′ junction for the transgenic and wild-type alleles, respectively. **(B)** Representative agarose gel electrophoresis of PCR products amplified from genomic DNA of two heterozygous transgenic mice (lanes 1 and 2) and two wild-type mice (lanes 3 and 4). PCR products spanning the transgene-genome junctions were confirmed by Sanger sequencing. **(C)** Sequences of oligonucleotide primers used for PCR genotyping.

The assays were initially evaluated using genomic DNA isolated from ear-punch tissues from two Cre-positive and two Cre-negative mice. PCR products corresponding to the predicted transgenic and wild-type alleles were resolved by agarose gel electrophoresis. Products of the expected sizes were excised from the gels, purified, and subjected to Sanger sequencing. Sequence analysis confirmed the predicted transgene-genome junctions and the corresponding wild-type genomic locus (Fig. 2A). Primer sequences are shown in Fig.2C. These junction-specific PCR assays therefore provide a direct means of distinguishing the Nestin-CreERT2 transgenic allele from the endogenous wild-type locus and enable precise genotyping of this transgenic mouse line.

## Discussion

Here, we characterized the insertion site and genomic architecture of the Nestin-CreERT2 transgene at nucleotide resolution using long-read whole-genome sequencing. Tiling and multiple-sequence alignment of PacBio HiFi reads enabled reconstruction of the complete 13.7-kb transgene and its genomic junctions. The transgene contains approximately 5.8 kb of rat *Nes* 5′ regulatory sequence driving CreERT2, followed by downstream genomic elements of the rat *Nes* gene. The presence of restriction endonuclease sites flanking several components is consistent with assembly of the original construct using conventional molecular cloning techniques.

Transgene integration was accompanied by deletion of 17.4 kb of endogenous mouse genomic DNA. Such structural alterations are common consequences of random transgenesis. In a large-scale analysis of 40 widely used transgenic mouse lines, deletions and other structural variants ranging from a few base pairs to >1 Mb were identified at the integration site in more than half of the lines examined (12). Approximately one-third of the transgenes disrupted endogenous coding sequences through insertion into exons (12). Unexpected sequences, including undocumented cassettes and contaminating DNA fragments, were also frequently identified (12).

Consistent with these observations, the Nestin-CreERT2 transgene contains a 409-bp fragment with 100% sequence identity to the *Escherichia coli wecA* locus that is not derived from the known vector sequence. Bacterial genomic sequences unrelated to the cloning vector have previously been identified in approximately 25% of characterized transgene insertions and were proposed to result from contamination of DNA preparations used for microinjection (12). The functional significance of the *wecA* fragment in the Nestin-CreERT2 transgene is unknown; however, bacterial sequences associated with transgenes have been reported to contribute to transgene silencing (13,14).

The transgene integrated within intron 1 of *Kcnd2* in the reverse transcriptional orientation relative to the endogenous gene. *Kcnd2* spans approximately 514.3 kb, of which intron 1 comprises approximately 496.7 kb. Whether the 17.4-kb intronic deletion and/or presence of the transgene alters *Kcnd2* expression remains to be determined. Constitutive *Kcnd2* knockout mice are viable and do not exhibit an overt gross phenotype (15); nevertheless, loss of *Kcnd2*/Kv4.2 is associated with alterations in neuronal electrophysiology, seizure susceptibility, nociception, and hippocampal-dependent learning and memory (16–19). Thus, determining whether *Kcnd2* expression or function is altered in Nestin-CreERT2 mice may be relevant to the interpretation of phenotypes obtained using this line.

Finally, identification of both transgene-genome junctions enabled the design of allele-specific PCR assays spanning the 5′ and 3′ insertion sites. These assays distinguish the wild-type and transgenic alleles and therefore permit direct discrimination of heterozygous and homozygous transgenic animals. This overcomes an important limitation of conventional Cre genotyping, which establishes the presence or absence of the transgene but does not determine zygosity.

In conclusion, long-read whole-genome sequencing resolved the complete sequence and genomic integration site of the Nestin-CreERT2 transgene, revealed an associated 17.4-kb endogenous deletion and an unexpected bacterial DNA fragment, and enabled development of a PCR-based strategy for direct determination of transgene zygosity. These findings illustrate the utility of long-read sequencing for defining the molecular architecture of legacy transgenic mouse lines and improving their genetic characterization.

## Acknowledgments

This work was supported by NIH grants R01CA301569, R01CA271601, R01NS130209, R01NS139479 and R21NS139480 to A.C. During the preparation of this manuscript, the authors used the large language model ChatGPT (OpenAI) to assist with grammar, spelling, and sentence structure. The authors reviewed and edited all AI-generated revisions and take full responsibility for the content of the published article.

---

**Supplementary Figure 1.**
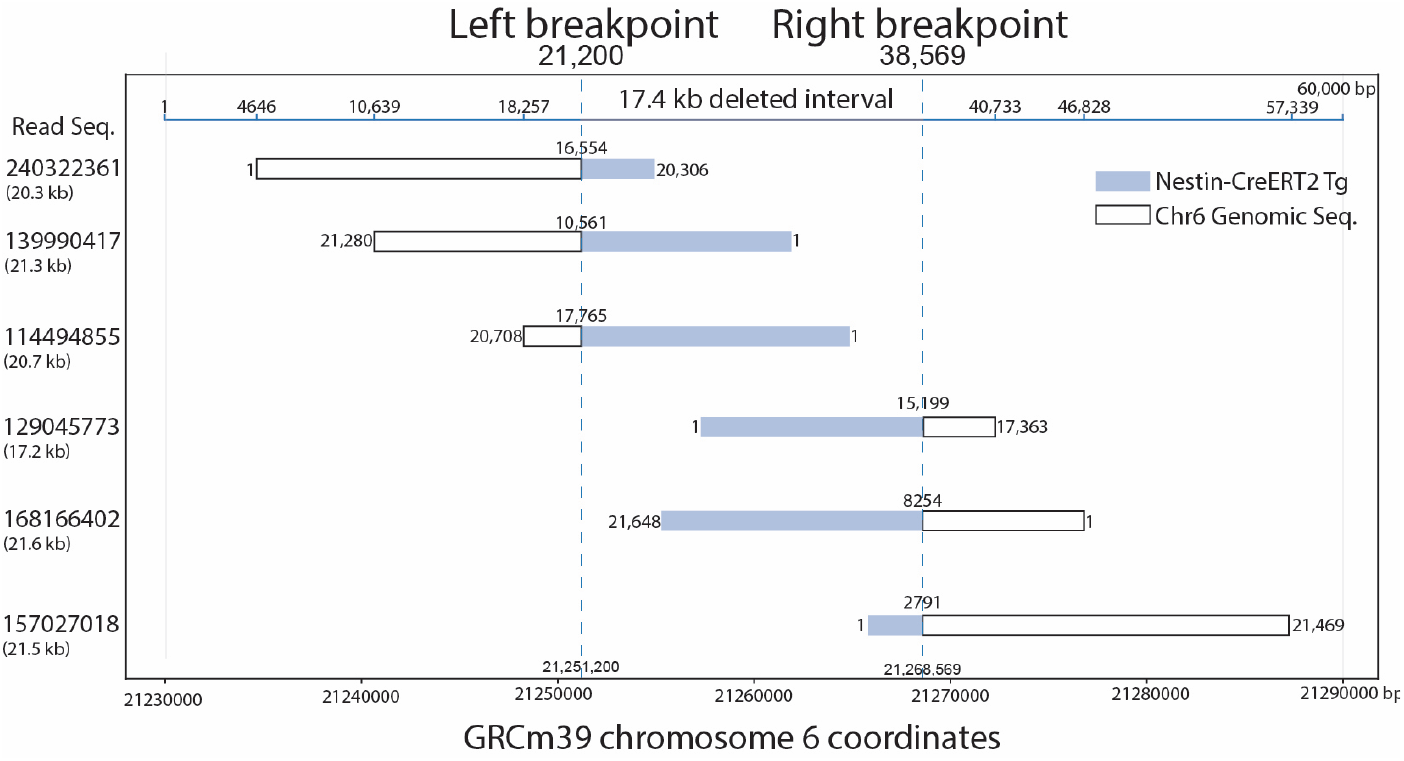
Supplementary PacBio HiFi reads spanning the Nestin-CreERT2 transgene integration junctions. Schematic representation of six PacBio HiFi reads supporting the transgene integration site, with three reads spanning the 5′ junction and three reads spanning the 3′ junction. Transgene-derived sequences are shown in blue and chromosome 6 genomic sequences in white. Dashed vertical lines indicate the genomic breakpoints at Chr6:21,251,200 and Chr6:21,268,569 (GRCm39), defining the approximately 17.4-kb deleted interval. PacBio HiFi read identifiers and read lengths are indicated on the left.

